# Uncovering large effect loci and epistasis for lifespan in standing genetic variation in the fruit fly *Drosophila melanogaster*

**DOI:** 10.1101/2024.10.05.616769

**Authors:** Joost van den Heuvel, Jelle Zandveld, Klaas Vrieling, Bart A. Pannebakker, Jan Kammenga, Bas J. Zwaan

**Affiliations:** Laboratory of Genetics, Wageningen University and Research, Droevendaalsesteeg 1, 6708 PB Wageningen, The Netherlands; Institute of Education Biology, Utrecht University, Padualaan 8, 3584 CH Utrecht, the Netherlands; Institute of Biology, Leiden University, Sylviusweg 72, P.O. Box 9505, 2300 RA Leiden, The Netherlands; Laboratory of Nematology, Wageningen University and Research, Droevendaalsesteeg 1, 6708 PB Wageningen, The Netherlands

**Keywords:** experimental evolution, pleiotropy, genetic variation, life history, ageing

## Abstract

Lifespan is a heritable trait with a polygenic architecture. Experimental evolution in combination with re-sequencing has often been used to identify candidate loci for lifespan in *Drosophila melanogaster*. Previous experiments showed that *Drosophila* populations experimentally evolved to increase late-life reproduction showed a correlated responses in development time, body size, but also lifespan. Subsequent whole genome sequencing allowed for the identification of candidate loci that correlated to lifespan differentiation. However, it remains difficult to assess whether candidate loci affect lifespan and to what extent such loci pleiotropically underpin multiple traits. Furthermore, recent studies indicate that lifespan effects of loci are often context dependent, but genotype-by-genotype interactions remain understudied. Therefore, here, we report on a study where we genotyped 3210 individuals for 32 candidate loci that emerged from our evolve and re-sequence experiment and tested, (1) whether these loci significantly affected lifespan, (2) the effect size of each locus, and, (3) how these loci mutually interact, i.e. determine the level of epistasis in moulding lifespan. Of the 32 loci, six showed significant main effect associations, of which three loci showed effects of 6.6 days difference in lifespan or larger, while the overall average lifespan was 41.7 days. Eight additional significant pairwise interactions between loci were found, of which four (single) main effects and one three-way interaction was significant. Lastly, we found that alleles that increased lifespan did not necessarily have higher frequencies in populations that showed increased lifespan, indicating that lifespan itself had not been the major target of selection. Our study indicates that individual genotyping following an evolve and re-sequencing study is essential to understand the mechanistic basis of polygenetic adaptation.

## Introduction

Understanding how genetic variation is generated, maintained, and altered in natural populations is a main theme in evolutionary biology. Of great interest is to explain the mechanistic genetic basis of the evolution of quantitative traits that are associated to fitness, such as survival. Lifespan, as a measure of survival, is a complex trait with a heritability ranging between 0.1 to 0.4 (Roff & Mousseau, 1987, Lehtovaara et al. 2013). In *Drosophila* several association studies using natural standing genetic variation have been performed to elucidate the genetic architecture of lifespan. Overall, three observations stand out from such studies. Firstly, while initially relatively low numbers of natural variants have been found, increased experimental and statistical power allowed for the identification of many more loci as candidates affecting lifespan (Huang et al. 2020, Pallares et al. 2022, Wong & Holman, 2023). Secondly, the effects of such variants typically are dependent on sex, environment, and other loci (Nuzhdin et al. 1997, Leips and Mackay 2002, Pallares et al, 2022) which in combination can show three-way interactions. Thirdly, only a small proportion of the phenotypic variance is explained by the uncovered allelic variation (Mackay 2002), as is the case for complex trait such as body size, for which many loci explain only a proportion of the phenotypic variation (Yang et al. 2010). Many complex traits seem to follow the theoretical expectation from Fisher’s infinitesimal model (Barton et al. 2017), which states that many loci of small effect underpin the quantitative variation of traits.

Lifespan is a life history trait for which trade-offs with fecundity are expected (Williams 1957, Kirkwood 1977), but also other traits are hypothesized to determine its phenotypic value in natural populations (Roff 1992, Stearns 1992). Furthermore, next to loci that underpin trade-offs, deleterious mutations can accumulate over generations, especially when expressed late in life where natural selection against such mutations is very weak (Medawar 1952). To test which other traits are genetically correlated to lifespan in *Drosophila*, selection for increased lifespan or postponed reproduction has often been employed in the past. This consistently led to an increased lifespan (Luckinbill et al. 1984, Rose 1984, Zwaan et al. 1995, Partridge et al. 1999, Stearns et al. 2000, May et al. 2019). In a number of these studies the genomes of the evolved populations were re-sequenced (Remolina et al. 2012, Carnes et al. 2015, Fabian et al. 2018, Hoedjes et al. 2019) which resulted in the identification of hundreds of loci that are candidates for lifespan regulation. However, these selection experiments also resulted in divergence for other traits, such as development time, body size, age-dependent fecundity and lifespan. Therefore, a key question remains whether the identified candidate loci in these evolve and re-sequence experiments directly affect lifespan.

To furher elucidate the genetic underpinning of lifespan of these candidate loci we associated variation at 32 candidate loci with lifespan in *Drosophila melanogaster*. Previously, we performed selection on late reproduction which resulted in increased lifespan, but also in development time, and body size (May et al. 2019) compared to populations selected on early reproduction (control populations). Using whole genome pooled resequencing we identified a high number of loci to respond to the two evolution treatments (Hoedjes et al. 2019). This previous analysis allows us to choose among 298 candidate loci that were consistently differentiated between early and late reproduction populations. This allowed us to sieve out candidates that are potentially causal to the lifespan differences observed between the diverged populations. By associating lifespan to the genotypes of 3210 individuals, we tested (1) whether they were signifcantly associated with lifespan, (2) how large their effects were and (3) how these loci mutually interacted in moulding lifespan. By measuring these effects we can make the first step to further understand genetic architecture of lifespan in natural populations, which is an important quantitative trait in the the evolution life histories.

## Method

### Fly populations, culture conditions

*Drosophila melanogaster* flies used in this study are part of a continuously selected set of populations that combines selection for differential larval food concentrations and age at reproduction (early or late) in a factorial design (May et al. 2019). A detailed description of the experimental design can be found in May et al. (2019). The six founding populations with which the experimental evolution was started were caught in a traject between Austria and Greece and crossed for 40 generations to allow for laboratory evolution and recombination between ancestral populations to decrease linkage disequilibrium (May et al. 2019). In the current experiment we focussed on loci that significantly differed in allele frequency between populations selected for early and late reproduction (Hoedjes et al. 2019). To maximize the probability of finding loci that affect lifespan in a population, we used individuals from one of the 24 selected populations, of which replicate #3 of the experimental regime “1x larval diet and early reproduction” (1-E3) was chosen. This regime was chosen as it can be considered the standard for laboratory propagation of *D. melanogaster* populations, and thus may serve as a control that can be compared to the late reproduction regime. Moreover, replicate #3 was chosen because when compared to other populations more candidate loci segregated at frequencies relatively close to 50% (see below for details), allowing for more even genotype distributions and thereby increasing the power of the study. Furthermore this allowed us to detect gene by gene interactions. Prior to the experiment, replicate #3 underwent the early reproduction experimental evolution treatment for 179 generations (May et al. 2019).

All experiments were performed at 25°C and 50% humidity on a 12-h light:12-h dark cycle. All flies were reared in standard density culture (42 µL of eggs, Clancy and Kennington 2001) on either 1x medium containing per litre 100 g sucrose, 70 g dried yeast (Fermipan): 20 g agar (Sigma), 15 ml nipagin and 3 ml proprionic acid, or 0.25x medium (25g sucrose, 17,5 g dried yeast, other components similar).

Prior to the experiment, flies were kept at 1x medium for two generations under standardized larval densities (42 µL of eggs, Clancy and Kennington 2001) removing potential maternal effects due to the selection regime. Then eggs were harvested on a petri dish with 1x medium. Initially, interactive effects between larval diet and genetic variation on lifespan were of interest to us as also a large evolutionary response was measured between flies evolved on different larval conditions. Therefore, a controlled number of eggs (42 µL of eggs, Clancy and Kennington 2001) were put on 0.25x and 1x diet in four replicates each in 65mL population bottles.

### Experimental setup

All flies were allowed to fully develop into adults and kept in mixed sex groups and emerged flies were kept in these conditions for at least a day to allow for mating. Then flies were pooled among four replicate bottles per larval diet, sedated with CO_2_ and female flies were randomly allotted to 34 half-pint bottles containing fresh (1x) medium 30 mL to a total of 100 females per bottle. After set up, flies were transferred to new bottles every two or three days. Dead flies were counted, removed, and individually frozen at -80°C until further analysis. Some flies escaped or were accidentally crushed during transfers from which no lifespan association could be made. In total we measured lifespan for 3210 individuals.

### Candidate loci

All flies in the lifespan assay were individually genotyped for 32 SNPs. These SNPs were chosen from our previous whole genome resequencing experiment (Hoedjes et al. 2019) based on the following criteria. In our previous analysis we identified 298 SNPs with significant allele frequency differences based on the GLM statistic (Hoedjes et al. 2019) and consistent differences between all early and late reproduction populations. In this paper, we calculated the average SNP allele frequency difference between populations selected for early and late reproduction. The candidate had to have at least a 0.3 difference in allele frequency (between 1x diet early reproduction and 1x late reproduction) resulting in 254 candidates. Secondly, SNPs were annotated whether they were found in coding regions and how many SNPs were found per gene. Genes with more than one significant SNPs were prioritized over those with only a single significant SNP. Thirdly, we used the variant annotation and prioritized according to non-synonymous > synonymous > intron variant / promotor region. If based on these criteria candidate loci had equal priority, those with the largest allele frequency differences were chosen. For each gene, only the highest differentiated SNP per gene was selected. This procedure resulted in 29 candidate SNPs (Table S1). This is therefore a 10% sample of all significant loci initially identified as those that consistently differ between early and late reproduction populations (Hoedjes et al. 2019). Additionally, three SNPs in the genes *hppy, Eip75B*, and *sick* were included in our analysis because these genes have been implicated to functionally affect life history traits including lifespan (Bryk et al. 2010, Lam et al. 2010, Huang et al. 2020, Parker et al. 2020, Hoedjes et al. 2023). However, the SNPs in these genes were less divergent in allele frequency than all other loci. The resulting list of candidates and their properties are listed in Table S1.

### DNA isolation, primer design, and KASP

To extract DNA, each sampled fly was put in a tube and grounded in 100 ul Squishing buffer (0.1 M Tris HCl pH 8, 1 mM EDTA, 0.25 M NaCl, 1 µg/ul Proteinase K), incubated for 45 minutes at 55°C followed by two minutes at 95°C to inactivate Proteinase K. The extract was centrifuged for three minutes at 13000 rpm and after centrifugation 50ul of the supernatant was transferred to a clean tube. The DNA concentration of each sample was normalized to 5 ng/ul per sample. Samples were stored in the freezer until use. All samples were genotyped using a Kompetitive Allele Specific PCR (KASP) genotyping assay (Semagn et al. 2014). In a KASP assay two allele-specific forward primers are designed that carry a tail which is labelled with a dye, FAM or HEX, by which for each candidate locus the genotype can be detected for each individual by relative dye detection in a qPCR machine. For 32 SNP loci, KASP primers were designed using the Kraken software (http://www.lgcgenomics.com) and ordered at Integrated DNA Technologies (Supplement File 1).

DNAs were further diluted to 1 ng/µL and analyzed on the LGC genomics SNP genotyping line according the manufactures’ instructions. Genotypes were automatically called using the Kraken software and manually adapted when necessary (Semagn et al. 2014).

### Statistical analysis

All analysis were performed using the R software (v3.6.1, R core Team 2019).

#### Population genetics

First, we tested whether genotype frequencies deviated from Hardy-Weinberg equilibrium with a Chi-square test. Furthermore, we estimated the linkage disequilibrium between all pairwise SNPs using the *LD() function* from the genetics package (Warnes 2003). We used the r^2^ statistic for LD and the p-value to test whether this statistic was significantly different from 0. The p-values were adjusted for multiple testing using *p*.*adjust()* with the ‘BH’ method (Benjamini and Hochberg 1995).

#### Main effects

We fitted a linear mixed effects model for lifespan with genotypes of a particular SNP and larval condition as main effects and replicate bottle as random effect using *lmer* (lme4 package, Bates et al. 2015) using a normal distribution as error structure. Then we fitted a model without genotype effects. The difference between these models indicates the significance of genotype for each locus, and the significance of this difference was tested using the *anova()* method, using ‘chisq-test’. The p-values were adjusted for multiple testing using *p*.*adjust()* with the ‘BH’ method (Benjamini and Hochberg 1995). All adjusted p-values below 0.05 were considered significant. The variance explained was calculated using the r.squaredGLMM(model) function of the (MuMIn package, v 1.46.0, Barton 2022) where ‘model’ was an above-described *lmer* model containing only the fixed effects for which the variance needed to be explained.

#### Test for genotype-by-genotype interactions

To test for genotype-by-genotype interactions we fitted a model with lifespan as the dependent variable and genotype effects of SNP1, genotype of SNP2, and larval condition as main effects, including interaction (I1). Next to this, we fitted a model without interaction (I2). Similarly, as with the main effects, we tested for the differences using the anova() method, using ‘chisq-test’. The p-values were adjusted for multiple testing using p.adjust() with the ‘BH’ method (Benjamini and Hochberg 1995). All adjusted p-values below 0.05 were considered significant. We only tested for interactions between genotypes when a minimum of five individuals was counted for each pairwise genotype combination for both genes in each larval condition. In total, this led to 366 tests. Similarly, we tested all three-way interactions for which at least five individuals were present for each multilocus genotype, amounting to a total of 1758 tests. For three-way interactions, the models with three-way interactions were compared to the models with three two-way interactions, and p values were adjusted as described above.

## Results

### Allele frequencies, HW equilibrium

The minor allele frequency of only one SNP was below 0.1 and for three additional SNPs between 0.1 – 0.2. Six loci showed a significant deviation from Hardy Weinberg equilibrium (p.adj<0.05, Table S2). The reference allele frequencies of the current study (generation 179) were significantly and highly correlated to those of the previous pooled resequencing experiment (Hoedjes et al. 2019) generation 115, Pearson r =0.87, t_30_=9.87, p<0.001, Figure S1), indicating that the allele frequencies from this study matched well with our previous whole genome sequencing experiment.

### Consistencies with whole genome sequencing

To study the similarity of our genotyping with our previous whole genome sequencing analysis, we calculated the correlation distance of allele frequencies for the 32 candidate genes with all other populations in the entire selection experiment (May et al. 2019). The current study was performed on the 1x diet early reproduction replicate #3 (1-E3). Using correlation distances of this study with all 24 populations, the allele frequencies of this study clustered most closely together with (1-E3) sequenced in generation 115 (Figure S1). This indicates not only that our genotyping has led to reproducible results but also that the population changes between these generations have been relatively small as compared to the differences between populations.

### Linkage disequilibrium

Linkage disequilibrium was estimated using the r^2^ statistic. As expected, r^2^ was higher for loci that were sampled in closer physical proximity (Figure S2). All combinations of SNPs that showed a r^2^ higher than 0.05 were found on the same chromosomal arms. Because 23 values of r^2^ were higher than 0.2, the linkage disequilibrium might be relevant for the independence of the association between genotypes and lifespan (Table S3).

### Main effects on lifespan

Not a single locus showed a significant interaction with larval diet. Flies reared on 1x diet had an average lifespan of 38.4 days, while flies reared on 0.25x diet had an increased lifespan of 6.8 days (Figure S3). Therefore, larval diet as main effect is considered in the statistical models.

From the 32 loci for which we tested the main effect of genotype, seven loci showed significant genotype effects on lifespan (p.adj<0.05, Table S4). However, as high levels of LD were found between some of the significant loci, we tested whether the effects of these loci were independent. The model with *mRps22* and *CG9997* showed such a dependence, and after this analysis six independent loci remained significant (Figure 1), of which three loci showed very large effects. For instance, for the gene *hppy*, the lifespan difference between the shortest and longest living genotypes was on average 12.0 days. This was 7.4 days for *Eip75B* and 6.6 days for *mRps22*. The percentage of phenotypic variance explained by these variants alone was 10.3%, 6.4%, and 5.2% for *hppy, Eip75B* and *mRps22*, respectively (Table S5). For the three remaining loci, lifespan differences between genotypes fell between 1.1 and 2.3 days (Figure 1).

**Figure 1.**
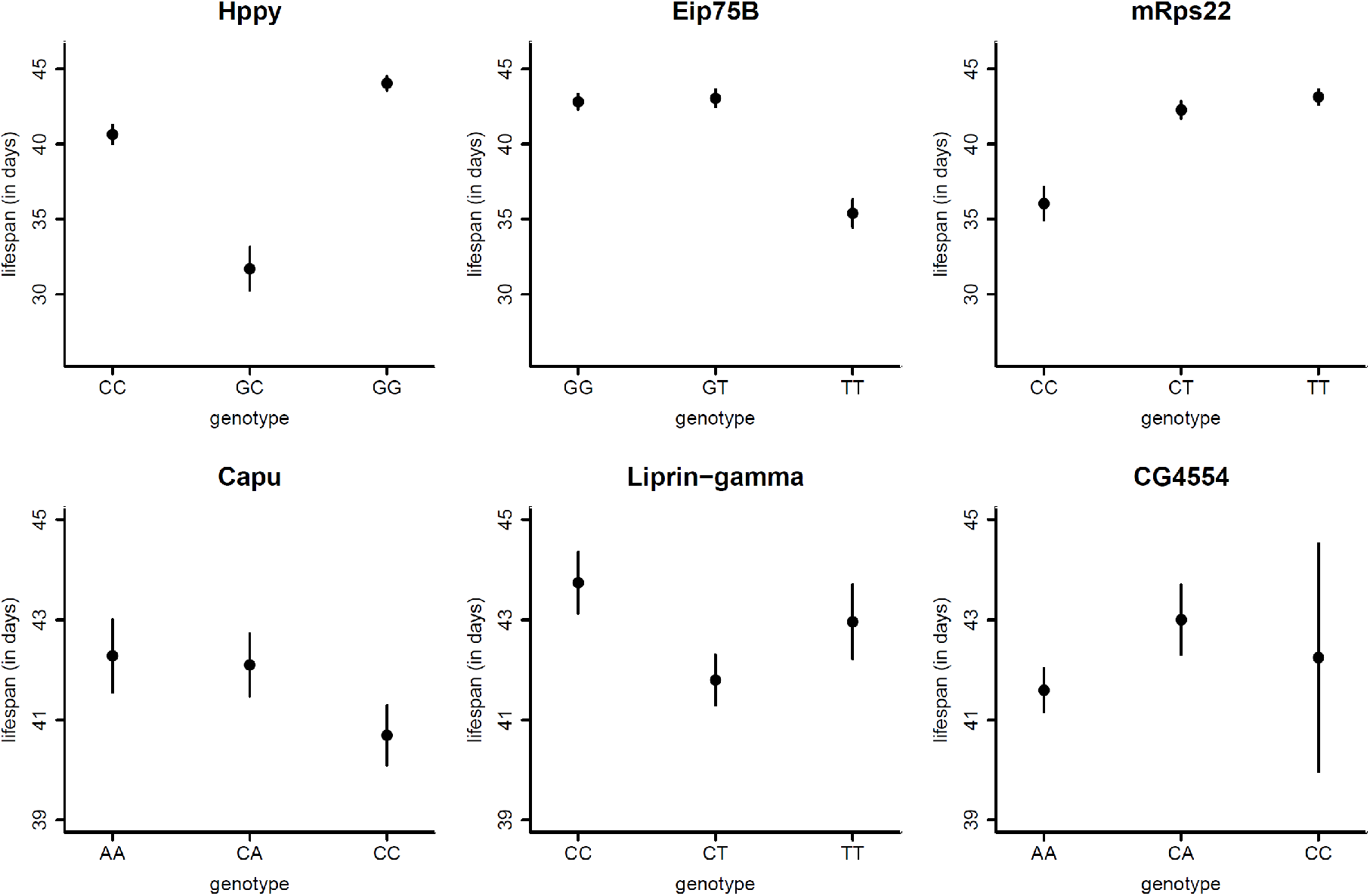
Mean and standard errors of lifespan (in days) dependent on genotypes for the six SNPs that tested significant for main effect associations. Titles of the subpanels indicate gene name. The x axis indicates the genotypes.

Interestingly, for both *Eip75B* and *mRps22*, one homozygote genotype stood out as having a relatively low lifespan, while the heterozygote had a similar lifespan as the other homozygote, indicating dominance for these alleles. For *hppy* the heterozygote was associated with the lowest lifespan, therefore showing underdominance. Furthermore, allele frequencies from five out of the six loci that significantly affected lifespan deviated from Hardy-Weinberg equilibrium, while from the other 26 loci (i.e. not significantly associated with a lifespan increase), only one showed a deviation from Hardy-Weinberg equilibrium. This result was significantly different from a random association (Fisher exact = 79.43, p<0.05). Thus, loci that are significantly associated with lifespan were more likely to be out of Hardy-Weinberg equilibrium.

### Gene-by-gene interaction

We tested whether the pairwise interactions between loci were significant (as compared to both main effects). Indeed, eight pairwise interactions were significant (i.e. taking the level of p.adj <0.05, Table 1). The three loci with the largest main effects, *hppy, Eip75B*, and *mRps22*, also showed significant interactions (Figure 2). Furthermore, four SNPs that did not affect lifespan as main effects showed a significant genotype-by-genotype interaction. For instance, lifespan effects of *mrfn* and *BOD1* interacted significantly with *mRps22*, and the effects of *su(var)3-7* and *CG33346* were dependent on the *capu* genotype (Figure 2). Therefore, testing for interactions increased the number of loci associated with lifespan. Furthermore, the addition of these interactive effects increased the explained phenotypic variation by 2.9% compared to when only main effects were considered. However, everysingle interaction had a relatively small effect, around or below 1% variance explained (Table 1). Lastly, one of the 1758 three-way interactions tested was significant. This was between loci on the genes *hppy, liprin-gamma*, and *Noa36* (Figure S4).

**Table 1.** Loci that significantly interact. The first two columns indicate the interacting loci, the 3^rd^, 4^th^ and 5^th^ column present the test statistic, raw and fdr adjusted p value and variance explained (R^2^).

| Locus A | Locus B | Chi<sup>2</sup> | Raw P value | Fdr adj. p value | Variance explained |
| --- | --- | --- | --- | --- | --- |
| Capu | CG33346 | 18.63 | 9.3*10<sup>-4</sup> | 0.044 | 0.57% |
| Capu | Su(var)3-7 | 24.99 | 5.1*10<sup>-5</sup> | 3.6*10<sup>-3</sup> | 0.74% |
| Hppy | Eip75B | 29.06 | 7.6*10<sup>-6</sup> | 1.2*10<sup>-3</sup> | 0.87% |
| Hppy | mRps22 | 20.47 | 4.0*10<sup>-4</sup> | 0.021 | 0.57% |
| Liprin-gamma | Eip75B | 26.33 | 2.7*10<sup>-5</sup> | 2.3*10<sup>-3</sup> | 0.78% |
| Eip75B | mRps22 | 23.00 | 1.3*10<sup>-4</sup> | 7.7*10<sup>-3</sup> | 0.64% |
| mRps22 | BOD-1 | 37.23 | 1.6*10<sup>-7</sup> | 6.8*10<sup>-5</sup> | 1.08% |
| mRps22 | Mfrn | 28.90 | 8.2*10<sup>-6</sup> | 1.2*10<sup>-3</sup> | 0.86% |

**Figure 2.**
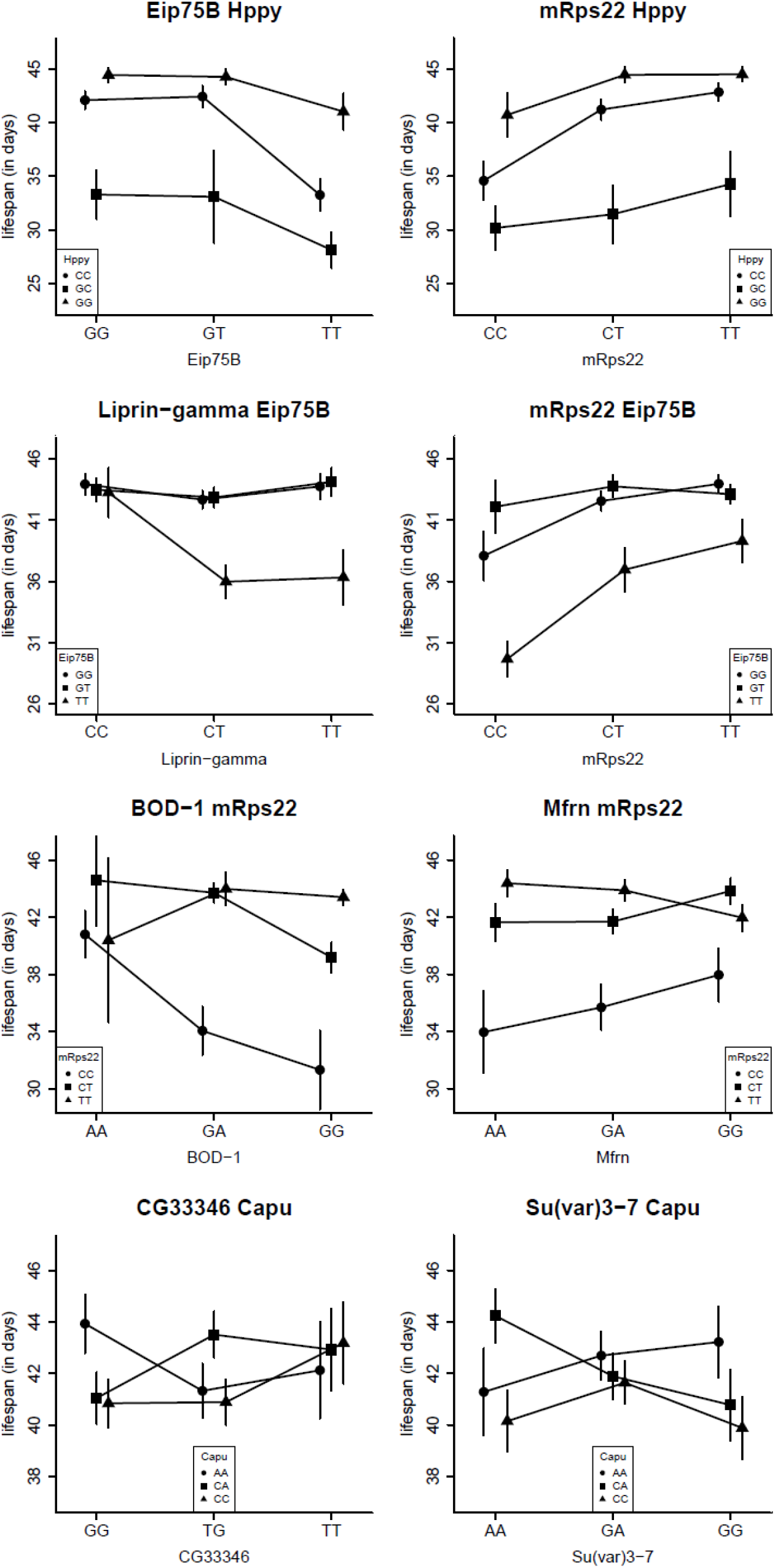
Combinations of loci for which interaction significantly affected lifespan (in days). Each panel title lists the gene names associated to the two interaction loci. The x axis and different symbols indicate the genotypes at the first and second gene respectively mentioned in the title of each subpanel. Error bars indicate standard errors on the mean.

### Direction of lifespan effects

Our experimental evolution populations were selected on the ability to reproduce late in life, with early reproduction serving as a control, as this is the common fly-maintenance practice in most fly laboratories. As a correlated response, lifespan was increased in late reproduction populations (May et al. 2019). If selection on the experimental evolution had acted on alleles that only increased lifespan, it would be expected that these alleles would have increased in frequency in late reproduction populations. We would therefore expect that alleles that positively associate with lifespan in this experiment would have an increased allele frequency in late reproduction populations. Given the result shown in Figure 1, we expect the longer-lived populations to have a decreased frequency of the C, T, C, C, T, and A alleles for the genes *hppy, Eip75B, mRps22, capu, Liprin-gamma*, and *CG4554*, respectively. However, this expectation is only born out for *hppy* and *CG4554*, thus the minority of the loci (Figure S5). This indicates that direct selection on lifespan may not causally underpin the allelic differentiation between the populations selected for early and late reproduction.

## Discussion

Selection for postponed reproduction in an outbred population of *Drosophila melanogaster* resulted in extended lifespan (May et al. 2019) and subsequently, by using whole genome sequencing, candidate loci for lifespan have been identified (Hoedjes et al. 2019). Here, we report on a follow-up study of this ‘evolve and re-sequence’ project. We tested 32 candidate loci for their effects on lifespan in a large cohort (n=3210). From the 32 tested loci, we found 11 loci that were significantly associated with lifespan. With an average lifespan of 41.7 days over all genotypes and conditions combined, we identified three loci to show very large effects on lifespan (between 6.6 and 12 days), while for three additional loci we found significant albeit smaller effects (i.e. between 1.1 and 2.3 days). Unexpectedly, when fitted together in one model, the three loci with the largest effect explained together 15.5% of the phenotypic variation. On top of the main effects, an additional four loci were identified through pairwise interactions. Furthermore, a last locus was significant in a three-way interaction without showing a significant main effect or a two-way interaction. These results show that following up a ‘evolve and re-sequence’ project with a candidate gene approach is a successful method for identifying whether candidate loci are involved in lifespan regulation in a population harbouring natural variation. On the flip side, this also means that the other 21 loci were not attributed to variation in lifespan. This could result from too low power to detect small effects on lifespan. Alternatively, these loci could be involved in the regulation of other traits, such as fecundity, development time, and body size, which are also differentiated between early and late reproduction populations (May et al., 2019).

To understand how organisms evolve, knowledge of the genetic architecture of the traits under selection is critical. How many variants segregate in a population and their effects remains elusive for many traits. Predicting which variants respond to selection is difficult and becomes even harder when many loci segregate with small effects or when interactions between loci occur (epistasis). Although we found large effect loci (making it easier to predict what the genetic course of evolution would be), we also found epistasis between 10 of the 32 candidate loci. While the variance explained by these effects was often small, it increases the complexity of the architecture of traits that has consequences for the predictability of evolution. Indeed, as recently shown, explicitly addressing epistasis improves our understanding of the genetics of ageing and diseases (Kiel et al. 2021, Wang et al. 2021).

We find large effect loci and interactions that contribute to lifespan variation, which is in line with two recent studies. Huang et al. (2020) previously estimated that the effects of some loci on lifespan can be large, similar to our findings, although the significance in that study only reached a ‘nominal’ level (of 10^−5^). Furthermore, genotype effects on lifespan are context-dependent, namely with temperature (Huang et al. 2020) and diet (Pallares et al. 2022). This context dependence of the expression of single loci can be extended to genotype-by-genotype interactions, of which we found ample evidence. Ten out of 11 loci showed significant interactions, and 2.9% of the total variance can be attributed to all two-way interactions. If interactions at the SNP level are important, we would expect relatively low overlap between SNP candidates in evolve and re-sequence experiments. Indeed, both Fabian et al. (2018) and Hoedjes et al. (2019) reported significant overlap between genes when comparing the results of similar evolve and re-sequence studies, but not at the level of SNPs. Although currently beyond the scope of this paper, in future studies it should be addressed whether the main and interactive effects are also present in other experimental evolution populations.

Some of the significant SNPs were found in genes that have been functionally tested for lifespan effects before. For instance, recently, it was found that RNAi on *Eip75B, CG4554*, and *capu* altered lifespan and age-specific fecundity (Parker et al. 2020, Hoedjes et al. 2023, Huang et al. 2020). Genome editing of the exact similar nucleotide studied in this experiment resulted in age-dependent effects on fecundity of *Eip75B*, but without a lifespan effect (Hoedjes et al. 2023). The large effect on lifespan in our study was dependent on the genetic background of another locus, namely, *liprin-gamma* (Figure 2) and therefore it is expected that the effects of genome editing is also context-dependent. It is well described that *hppy* interacts with the mTor-signaling pathway regulating life-history traits and apoptosis (Bryk et al. 2010, Lam et al. 2010) and has been previously identified in a QTL screening of lifespan (Stanley et al. 2017). It is therefore not surprising to find this locus associated with lifespan. While some of the studied genes are private to insects, others are found in a wider range of organisms. For instance, mutations in *mRps22* are known to cause several types of disorders in humans (Saada et al. 2007, Smits et al. 2011, Chen et al. 2018). In this study, we found a double non-synonymous mutation in *mRps22* to be associated with lifespan with a relatively high frequency. Other genes *liprin-gamma, su(var)3-7, BOD1, mfrn, CG33346* and *Noa36* have not yet been identified for their lifespan effects and can therefore be considered new candidates for further lifespan studies.

Evolutionary explanations of ageing are either based on deleterious mutations (mutation accumulation, Medawar 1952) or trade-offs between lifespan and other traits such as fecundity (antagonistic pleiotropy; Williams 1957, Kirkwood 1977). Interestingly, if mutation accumulation would explain most of the variation in ageing rate and lifespan, one would expect that alleles with higher frequencies in long-lived populationss have a positive effect on lifespan. Conversely, alleles that have increased in frequency in late reproduction populationss should also have been associated with higher lifespan. In four out of the six significant main effect SNPs, this was not the case. Hence, it is more likely that these loci have changed in frequency between the early and late-reproducing populationss because they are associated with other traits that are under selection. Indeed, our selection regime, that of reproduction at 14 (early reproduction) or 28 (late reproduction) days after egg laying is unlikely to result in a large selection differential directly on lifespan; most flies die long after these 28 days after egg laying of late reproduction.

Some of the genes that are included in this study have been functionally analyzed and this indicated that more traits than lifespan alone are regulated by these genes (Parker et al. 2020, Hoedjes et al. 2023). This corresponds to lifespan being a life history trait, trading off with other traits (Roff 1992, Stearns 1992). While the evolve and resequencing approach is clearly a powerful tool for identifying potential causal loci segregating in natural populations, our study indicates that it remains important to identify which traits are associated to genetic variation. Our future work will focus on assessing whether loci under selection affect other traits such as development time, body size, and age-dependent reproduction, as life history theory suggests.

## Supplemental figures and tables

**Figure S1.**
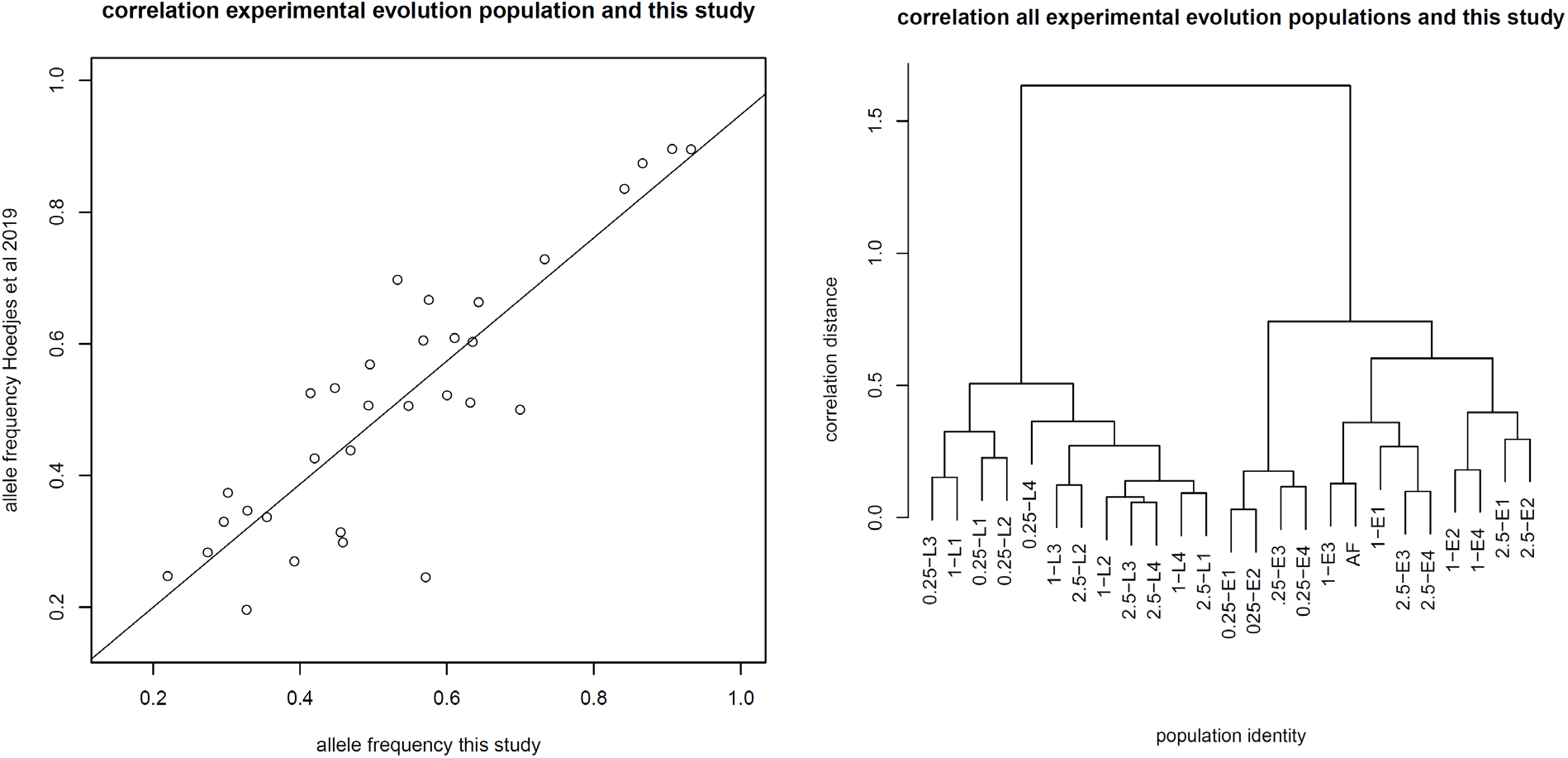
Correlation of allele frequencies of the 32 assayed loci between the pool-seq experiment (generation 115 of population CE3) and the KASP analysis (generation 179 of population 1-E3). (A) Correlation between the KASP analysis from this study and the pool-seq experiment. Line indicates fit of linear model (ANOVA) with a slope of 0.935 (p<0.001) and a non-significant intersect of 0.013. (B) Cluster dendrogram (Hierarchical clustering using correlation distance [d=1-|r|], where |r| is the absolute Pearson correlation coefficient) of all pool-seq re-sequenced populations and the allele frequencies of the KASP analysis (indicated by AF).

**Figure S2.**
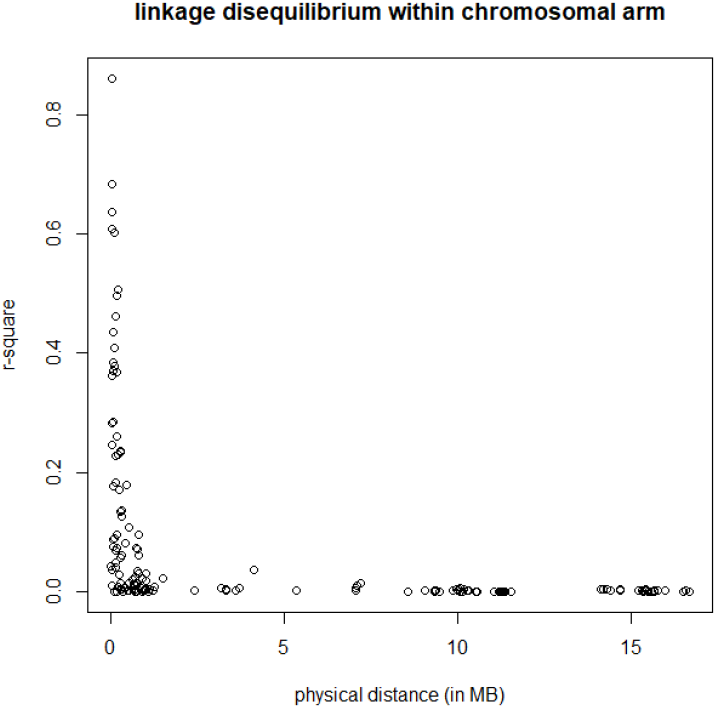
Linkage disequilibrium (r^2^) dependent on physical distance (x axis) for all possible combinations of loci within all a chromosomal arm.

**Figure S3.**
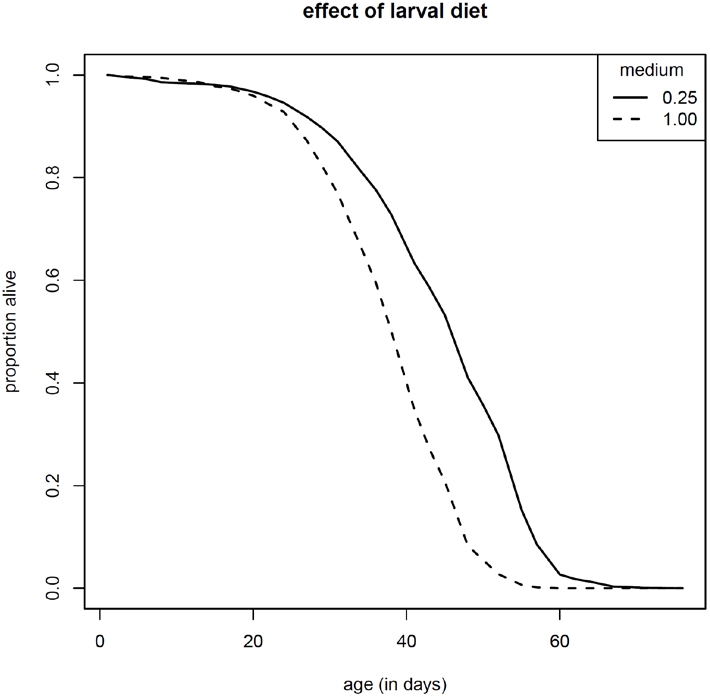
Survival curves of all individuals on 0.25x (solid line) or 1x larval diet (dashed line). The average difference between 0.25x and 1x

**Figure S4.**
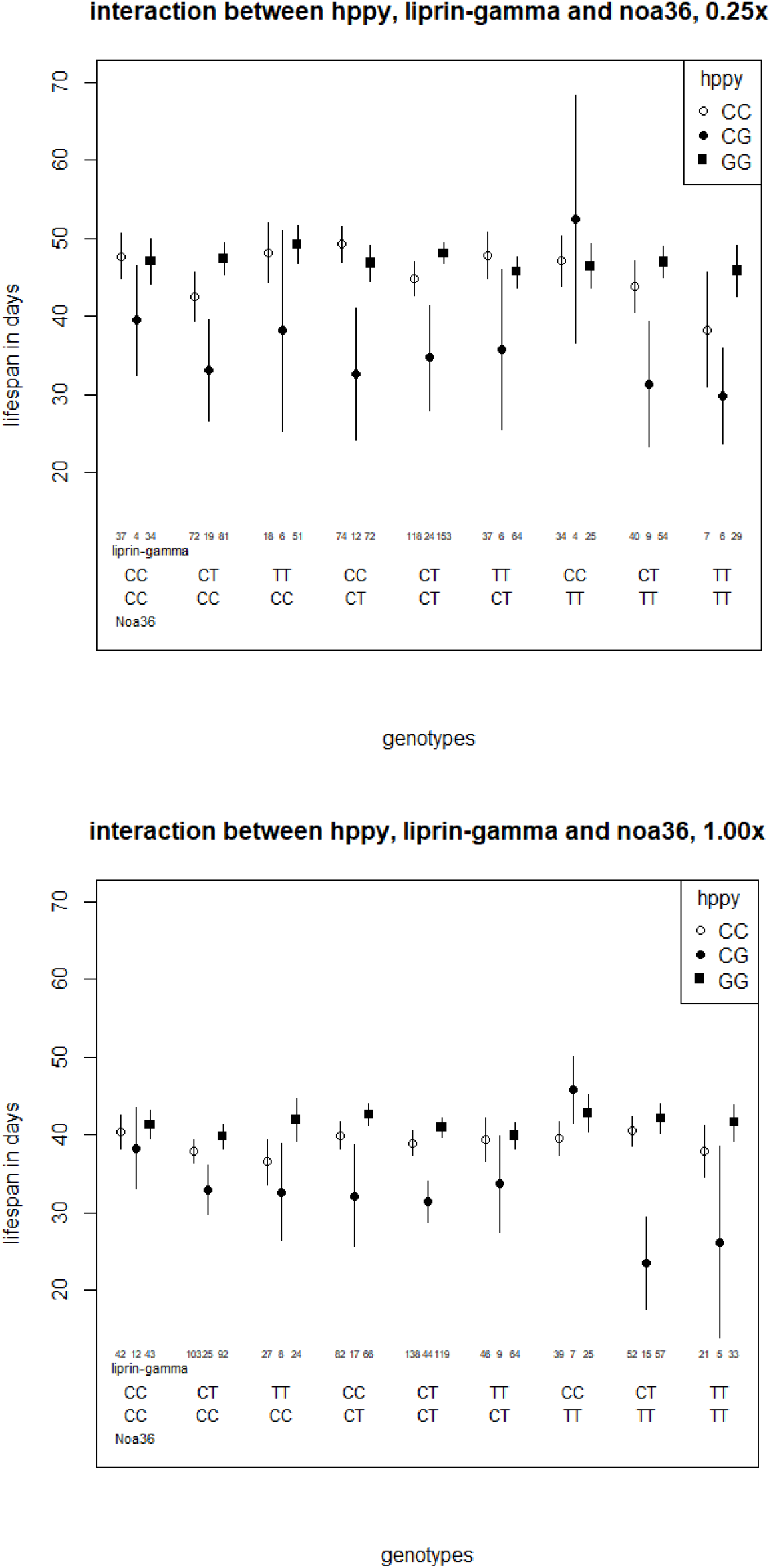
Three way interaction between hppy, liprin-gamma and Noa36. Top figure shows lifespan deendent on the genetic variation in these three genes for 0.25 larval diet, the bottom for 1.00 larval diet. Numbers indicate number of individual carrying a specific genotype.

**Figure S5.**
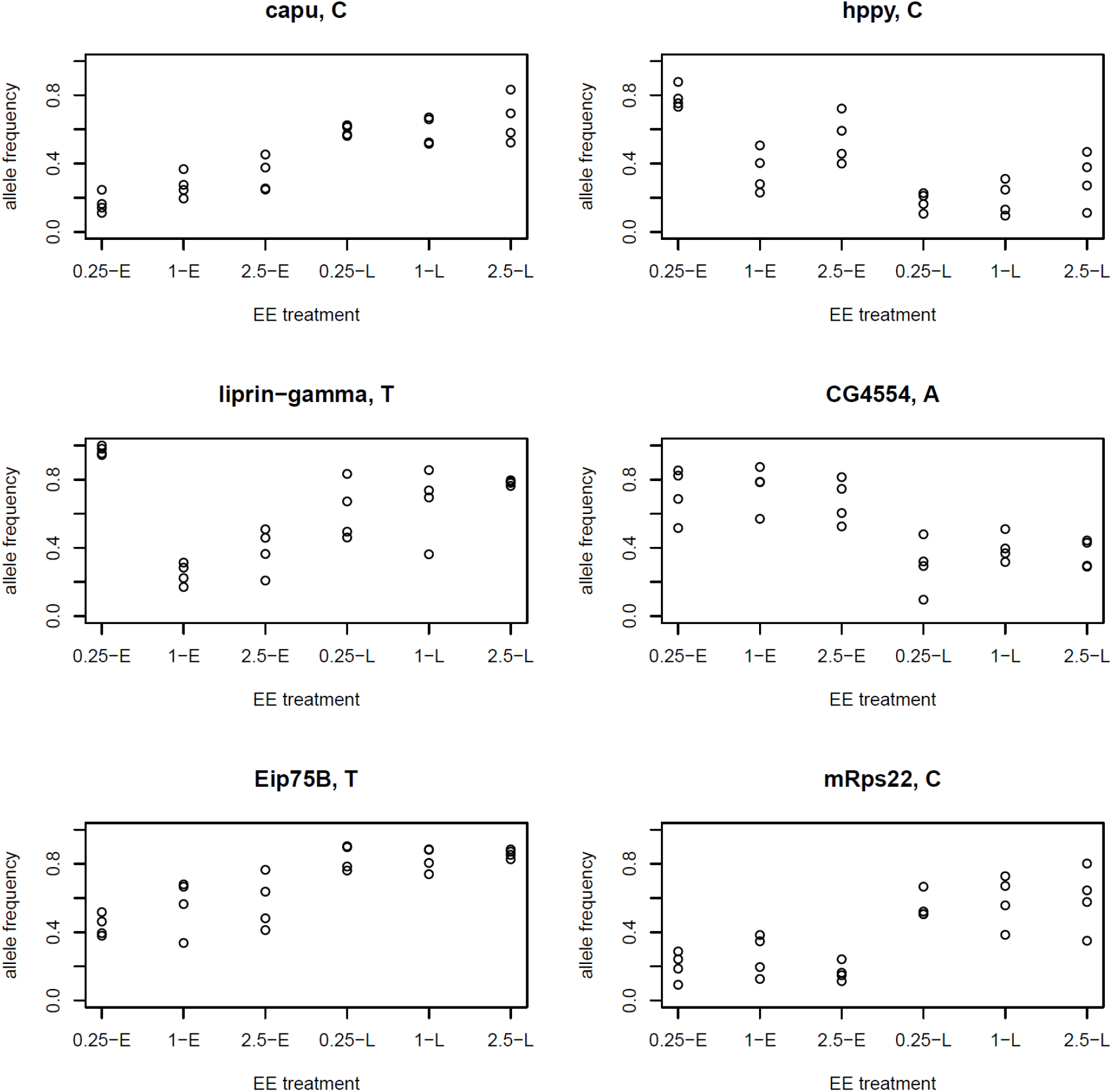
Allele frequency of reference alleles for the 6 main effects loci, as dependent on experimental evolution population (x axis). The treatment levels of the experimental evolution populationss differ in larval diet (0.25x, 1x and 2.5x) and adult selection for reproduction (early = E, and late = L). The alleles chosen to depict were those that associated with decreased lifespan and hence, were expected to be those that have higher frequency in the early reproduction populations (0.25-E, 1-E and 2.5-E). This was only the case for CG4554 and hppy.

**Table S1.**
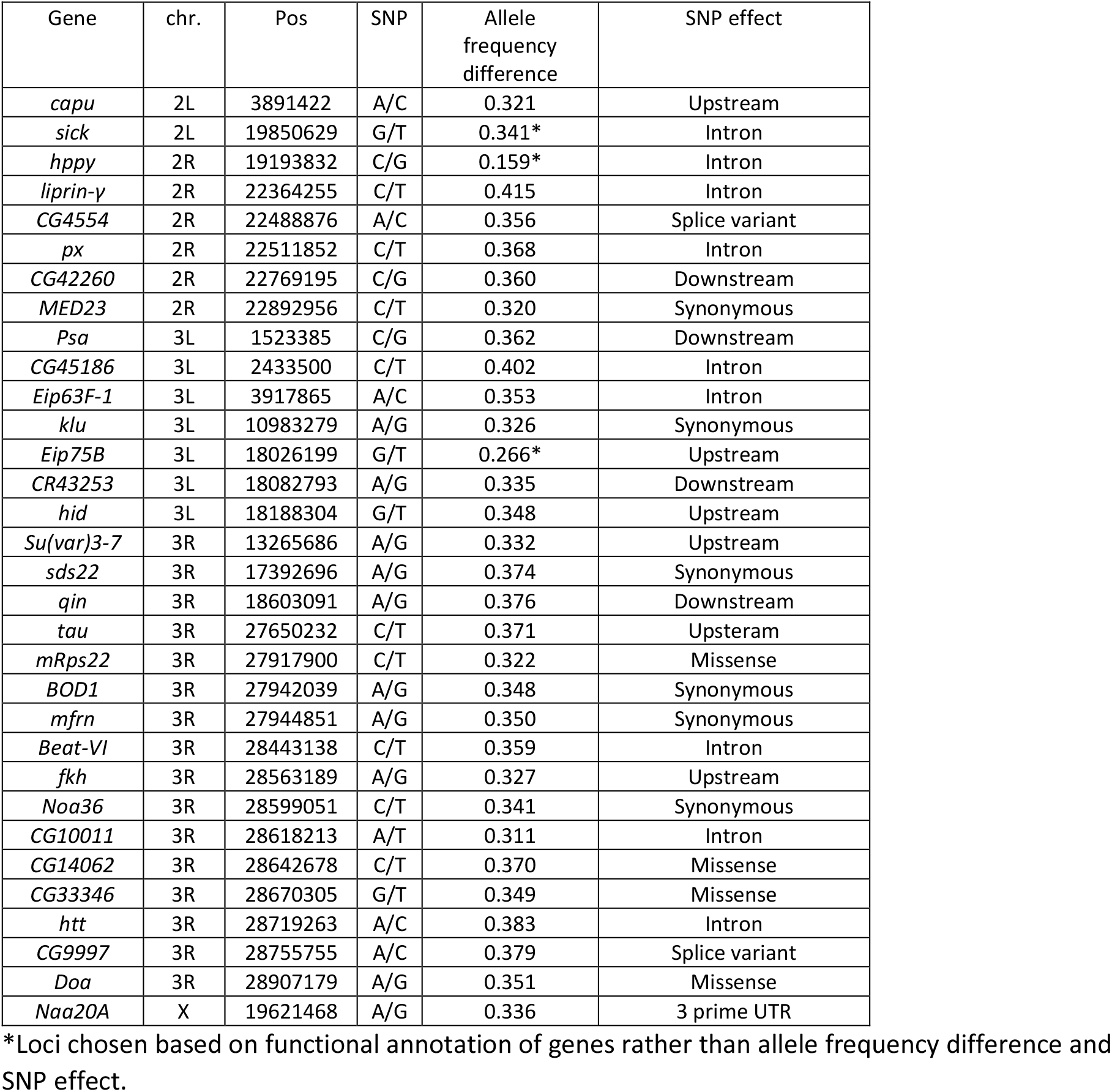
Variant information on which the candidates were chosen. The fifth and sixth columns indicate the allele frequency difference between early and late populations and the effect of the SNP.

| Gene | chr. | Pos | SNP | Allele frequency difference | SNP effect |
| --- | --- | --- | --- | --- | --- |
| *capu* | 2L | 3891422 | A/C | 0.321 | Upstream |
| *sick* | 2L | 19850629 | G/T | 0.341* | Intron |
| *hppy* | 2R | 19193832 | C/G | 0.159* | Intron |
| *liprin-γ* | 2R | 22364255 | C/T | 0.415 | Intron |
| *CG4554* | 2R | 22488876 | A/C | 0.356 | Splice variant |
| *px* | 2R | 22511852 | C/T | 0.368 | Intron |
| *CG42260* | 2R | 22769195 | C/G | 0.360 | Downstream |
| *MED23* | 2R | 22892956 | C/T | 0.320 | Synonymous |
| *Psa* | 3L | 1523385 | C/G | 0.362 | Downstream |
| *CG45186* | 3L | 2433500 | C/T | 0.402 | Intron |
| *Eip63F-1* | 3L | 3917865 | A/C | 0.353 | Intron |
| *klu* | 3L | 10983279 | A/G | 0.326 | Synonymous |
| *Eip75B* | 3L | 18026199 | G/T | 0.266* | Upstream |
| *CR43253* | 3L | 18082793 | A/G | 0.335 | Downstream |
| *hid* | 3L | 18188304 | G/T | 0.348 | Upstream |
| *Su(var)3-7* | 3R | 13265686 | A/G | 0.332 | Upstream |
| *sds22* | 3R | 17392696 | A/G | 0.374 | Synonymous |
| *qin* | 3R | 18603091 | A/G | 0.376 | Downstream |
| *tau* | 3R | 27650232 | C/T | 0.371 | Upstream |
| *mRps22* | 3R | 27917900 | C/T | 0.322 | Missense |
| *BOD1* | 3R | 27942039 | A/G | 0.348 | Synonymous |
| *mfrn* | 3R | 27944851 | A/G | 0.350 | Synonymous |
| *Beat-VI* | 3R | 28443138 | C/T | 0.359 | Intron |
| *fkf* | 3R | 28563189 | A/G | 0.327 | Upstream |
| *Noa36* | 3R | 28599051 | C/T | 0.341 | Synonymous |
| *CG10011* | 3R | 28618213 | A/T | 0.311 | Intron |
| *CG14062* | 3R | 28642678 | C/T | 0.370 | Missense |
| *CG33346* | 3R | 28670305 | G/T | 0.349 | Missense |
| *htt* | 3R | 28719263 | A/C | 0.383 | Intron |
| *CG9997* | 3R | 28755755 | A/C | 0.379 | Splice variant |
| *Doa* | 3R | 28907179 | A/G | 0.351 | Missense |
| *Naa20A* | X | 19621468 | A/G | 0.336 | 3 prime UTR |

\*Loci chosen based on functional annotation of genes rather than allele frequency difference and SNP effect.

**Table S2.** Overview of candidate genes characteristics. Gene name, chromosomal arm and position on it, SNP variation and counts for 0.25x diet and 1x diet genotypes are given. In the first column (capu) SNP variation (A/C) is indicated. Column 1/1 indicates count for A/A, 1/2 for AC and 2/2 for C/C. In the first from last column minor allele frequency (MAF) is given. The last column indicates whether all the genotype counts (A), those from 0.25x diet reared flies and / or those from the 1x diet reared flies are significantly different from Hardy-Weinberg equilibrium (p<0.05).

| Gene | chr. | Pos | SNP | 0.25x diet |  |  | 1x diet |  |  | MAF | HWE |
| --- | --- | --- | --- | --- | --- | --- | --- | --- | --- | --- | --- |
|  |  |  |  | 1/1 | 1/2 | 2/2 | 1/1 | 1/2 | 2/2 |  |  |
| *capu* | 2L | 3891422 | A/C | 381 | 480 | 551 | 385 | 525 | 636 | 0.4288 | A/0.25/1 |
| *sick* | 2L | 19850629 | G/T | 375 | 652 | 308 | 434 | 714 | 326 | 0.4689 |  |
| *hppy* | 2R | 19193832 | C/G | 476 | 110 | 587 | 619 | 173 | 542 | 0.4932 | A/0.25/1 |
| *liprin-γ* | 2R | 22364255 | C/T | 412 | 785 | 288 | 426 | 811 | 282 | 0.4554 |  |
| *CG4554* | 2R | 22488876 | A/C | 1056 | 309 | 26 | 1118 | 339 | 35 | 0.1335 | A/1 |
| *px* | 2R | 22511852 | C/T | 10 | 264 | 1259 | 17 | 267 | 1320 | 0.0932 |  |
| *CG42260* | 2R | 22769195 | C/G | 1296 | 164 | 8 | 1360 | 210 | 11 | 0.0676 |  |
| *MED23* | 2R | 22892956 | C/T | 923 | 500 | 69 | 950 | 535 | 87 | 0.2198 |  |
| *Psa* | 3L | 1523385 | C/G | 451 | 618 | 218 | 468 | 708 | 264 | 0.4199 |  |
| *CG45186* | 3L | 2433500 | C/T | 580 | 718 | 211 | 662 | 716 | 213 | 0.3681 |  |
| *Eip63F-1* | 3L | 3917865 | A/C | 699 | 640 | 134 | 766 | 676 | 126 | 0.3019 |  |
| *klu* | 3L | 10983279 | A/G | 209 | 671 | 586 | 188 | 737 | 626 | 0.3649 |  |
| *Eip75B* | 3L | 18026199 | G/T | 683 | 475 | 236 | 712 | 532 | 305 | 0.3549 | A/0.25/1 |
| *CR43253* | 3L | 18082793 | A/G | 232 | 696 | 555 | 217 | 784 | 569 | 0.3895 |  |
| *hid* | 3L | 18188304 | G/T | 698 | 541 | 86 | 777 | 606 | 115 | 0.2744 |  |
| *Su(var)3-7* | 3R | 13265686 | A/G | 366 | 750 | 359 | 418 | 778 | 396 | 0.4953 |  |
| *sds22* | 3R | 17392696 | A/G | 173 | 622 | 680 | 161 | 703 | 706 | 0.3273 |  |
| *qin* | 3R | 18603091 | A/G | 231 | 721 | 508 | 234 | 769 | 564 | 0.3997 |  |
| *tau* | 3R | 27650232 | C/T | 725 | 631 | 136 | 784 | 679 | 120 | 0.2963 |  |
| *mRps22* | 3R | 27917900 | C/T | 164 | 523 | 704 | 231 | 599 | 690 | 0.3284 | A/0.25/1 |
| *BOD1* | 3R | 27942039 | A/G | 104 | 558 | 789 | 117 | 601 | 831 | 0.2668 |  |
| *mfrn* | 3R | 27944851 | A/G | 282 | 754 | 403 | 303 | 738 | 494 | 0.4475 |  |
| *Beat-VI* | 3R | 28443138 | C/T | 132 | 573 | 730 | 140 | 682 | 739 | 0.3002 |  |
| *fkh* | 3R | 28563189 | A/G | 30 | 424 | 1066 | 29 | 444 | 1123 | 0.1582 |  |
| *Noa36* | 3R | 28599051 | C/T | 420 | 739 | 304 | 492 | 754 | 319 | 0.4523 |  |
| *CG10011* | 3R | 28618213 | A/T | 419 | 734 | 293 | 452 | 768 | 329 | 0.4584 |  |
| *CG14062* | 3R | 28642678 | C/T | 502 | 684 | 278 | 509 | 738 | 321 | 0.4321 |  |
| *CG33346* | 3R | 28670305 | G/T | 618 | 699 | 186 | 638 | 735 | 191 | 0.3567 |  |
| *htt* | 3R | 28719263 | A/C | 247 | 716 | 487 | 273 | 740 | 549 | 0.4143 |  |
| *CG9997* | 3R | 28755755 | A/C | 213 | 700 | 538 | 246 | 734 | 566 | 0.3924 |  |
| *Doa* | 3R | 28907179 | A/G | 280 | 750 | 491 | 283 | 772 | 542 | 0.4246 |  |
| *Naa20A* | X | 19621468 | A/G | 280 | 529 | 365 | 290 | 641 | 367 | 0.4672 | 0.25 |

**Table S3.** SNP combinations for which the linkage disequilibrium parameter r^2^ was found to be higher than 0.2. Columns list chromosomal arm, gene annotation, position of SNPs and exact r^2^.

| Chromosomal arm | geneA | geneB | Position SNP A on chromosome | Position SNP B on chromosome | $r^2$ |
| --- | --- | --- | --- | --- | --- |
| 2R | CG4554 | PX | 22488876 | 22511852 | 0.638 |
| 3L | Eip75B | CR43253 | 18026199 | 18082793 | 0.384 |
| 3L | Eip75B | hid | 18026199 | 18188304 | 0.602 |
| 3L | CR43253 | hid | 18082793 | 18188304 | 0.496 |
| 3R | mRpS22 | CG5514 | 27917900 | 27942039 | 0.363 |
| 3R | Beat.VI | CG10011 | 28443138 | 28618213 | 0.261 |
| 3R | Beat.VI | CG14062 | 28443138 | 28642678 | 0.230 |
| 3R | Noa36 | CG10011 | 28599051 | 28618213 | 0.683 |
| 3R | Noa36 | CG14062 | 28599051 | 28642678 | 0.610 |
| 3R | Noa36 | CG33346 | 28599051 | 28670305 | 0.368 |
| 3R | Noa36 | CG9997 | 28599051 | 28755755 | 0.228 |
| 3R | CG10011 | CG14062 | 28618213 | 28642678 | 0.861 |
| 3R | CG10011 | CG33346 | 28618213 | 28670305 | 0.461 |
| 3R | CG10011 | Htt | 28618213 | 28719263 | 0.285 |
| 3R | CG10011 | CG9997 | 28618213 | 28755755 | 0.378 |
| 3R | CG10011 | doa | 28618213 | 28907179 | 0.236 |
| 3R | CG14062 | CG33346 | 28642678 | 28670305 | 0.409 |
| 3R | CG14062 | Htt | 28642678 | 28719263 | 0.282 |
| 3R | CG14062 | CG9997 | 28642678 | 28755755 | 0.370 |
| 3R | CG14062 | doa | 28642678 | 28907179 | 0.233 |
| 3R | CG33346 | CG9997 | 28670305 | 28755755 | 0.246 |
| 3R | Htt | CG9997 | 28719263 | 28755755 | 0.436 |
| 3R | CG9997 | doa | 28755755 | 28907179 | 0.507 |

**Table S4.** Test statistics (chi), p values (P) and adjusted p values (Padj) for main effects and interaction with larval diet for SNP variation at 32 candidate loci.

|  |  |  |  | Main effect |  |  | Interaction |  |  |
| --- | --- | --- | --- | --- | --- | --- | --- | --- | --- |
| Gene | chr. | Pos | SNP | chi | P | Padj | chi | P | Padj |
| *capu* | 2L | 3891422 | A/C | 12.5896 | 0.0018 | 0.0118 | 0.3079 | 0.8573 | 0.9795 |
| *sick* | 2L | 19850629 | G/T | 2.2114 | 0.331 | 0.4814 | 0.0102 | 0.9949 | 0.9949 |
| *hppy* | 2R | 19193832 | C/G | 285.4659 | 1e-16 | 1e-16 | 15.4697 | 4e-04 | 0.014 |
| *liprin-γ* | 2R | 22364255 | C/T | 22.3943 | 1e-16 | 1e-04 | 0.4266 | 0.8079 | 0.9795 |
| *CG4554* | 2R | 22488876 | A/C | 11.4755 | 0.0032 | 0.0172 | 2.3887 | 0.3029 | 0.6462 |
| *px* | 2R | 22511852 | C/T | 4.3639 | 0.1128 | 0.2312 | 3.4871 | 0.1749 | 0.5739 |
| *CG42260* | 2R | 22769195 | C/G | 2.4206 | 0.2981 | 0.4567 | 0.2383 | 0.8877 | 0.9795 |
| *MED23* | 2R | 22892956 | C/T | 0.0692 | 0.966 | 0.966 | 1.7654 | 0.4137 | 0.7726 |
| *Psa* | 3L | 1523385 | C/G | 8.4891 | 0.0143 | 0.0574 | 0.6422 | 0.7253 | 0.9671 |
| *CG45186* | 3L | 2433500 | C/T | 0.2143 | 0.8984 | 0.9274 | 3.6739 | 0.1593 | 0.5739 |
| *Eip63F-1* | 3L | 3917865 | A/C | 0.2241 | 0.894 | 0.9274 | 4.0189 | 0.1341 | 0.5739 |
| *klu* | 3L | 10983279 | A/G | 0.9974 | 0.6073 | 0.7198 | 4.1964 | 0.1227 | 0.5739 |
| *Eip75B* | 3L | 18026199 | G/T | 228.6382 | 1e-16 | 1e-16 | 0.0579 | 0.9715 | 0.9949 |
| *CR43253* | 3L | 18082793 | A/G | 5.3967 | 0.0673 | 0.1795 | 2.4258 | 0.2973 | 0.6462 |
| *hid* | 3L | 18188304 | G/T | 2.9743 | 0.226 | 0.3807 | 0.8163 | 0.6649 | 0.9251 |
| *Su(var)3-7* | 3R | 13265686 | A/G | 4.6525 | 0.0977 | 0.2232 | 2.9337 | 0.2307 | 0.6151 |
| *sds22* | 3R | 17392696 | A/G | 1.5527 | 0.4601 | 0.5889 | 0.1575 | 0.9243 | 0.9859 |
| *qin* | 3R | 18603091 | A/G | 3.6486 | 0.1613 | 0.2868 | 0.2949 | 0.8629 | 0.9795 |
| *tau* | 3R | 27650232 | C/T | 7.8675 | 0.0196 | 0.0626 | 3.8728 | 0.1442 | 0.5739 |
| *mRps22* | 3R | 27917900 | C/T | 133.4492 | 1e-16 | 1e-16 | 7.947 | 0.0188 | 0.3009 |
| *BOD1* | 3R | 27942039 | A/G | 1.1068 | 0.575 | 0.7077 | 1.065 | 0.5871 | 0.854 |
| *mfrn* | 3R | 27944851 | A/G | 2.41 | 0.2997 | 0.4567 | 1.6667 | 0.4346 | 0.7726 |
| *Beat-VI* | 3R | 28443138 | C/T | 1.9144 | 0.384 | 0.5342 | 2.7199 | 0.2567 | 0.6318 |
| *fkf* | 3R | 28563189 | A/G | 0.8744 | 0.6458 | 0.7381 | 3.2463 | 0.1973 | 0.5739 |
| *Noa36* | 3R | 28599051 | C/T | 3.702 | 0.1571 | 0.2868 | 4.1639 | 0.1247 | 0.5739 |
| *CG10011* | 3R | 28618213 | A/T | 4.7527 | 0.0929 | 0.2232 | 3.3414 | 0.1881 | 0.5739 |
| *CG14062* | 3R | 28642678 | C/T | 6.9974 | 0.0302 | 0.088 | 3.8807 | 0.1437 | 0.5739 |
| *CG33346* | 3R | 28670305 | G/T | 4.3152 | 0.1156 | 0.2312 | 1.9041 | 0.386 | 0.7719 |
| *htt* | 3R | 28719263 | A/C | 1.6263 | 0.4435 | 0.5889 | 1.537 | 0.4637 | 0.781 |
| *CG9997* | 3R | 28755755 | A/C | 9.457 | 0.0088 | 0.0404 | 1.1173 | 0.572 | 0.854 |
| *Doa* | 3R | 28907179 | A/G | 7.9487 | 0.0188 | 0.0626 | 1.2244 | 0.5421 | 0.854 |
| *Naa20A* | X | 19621468 | A/G | 0.4377 | 0.8035 | 0.8866 | 0.312 | 0.8556 | 0.9795 |

**Table S5.**
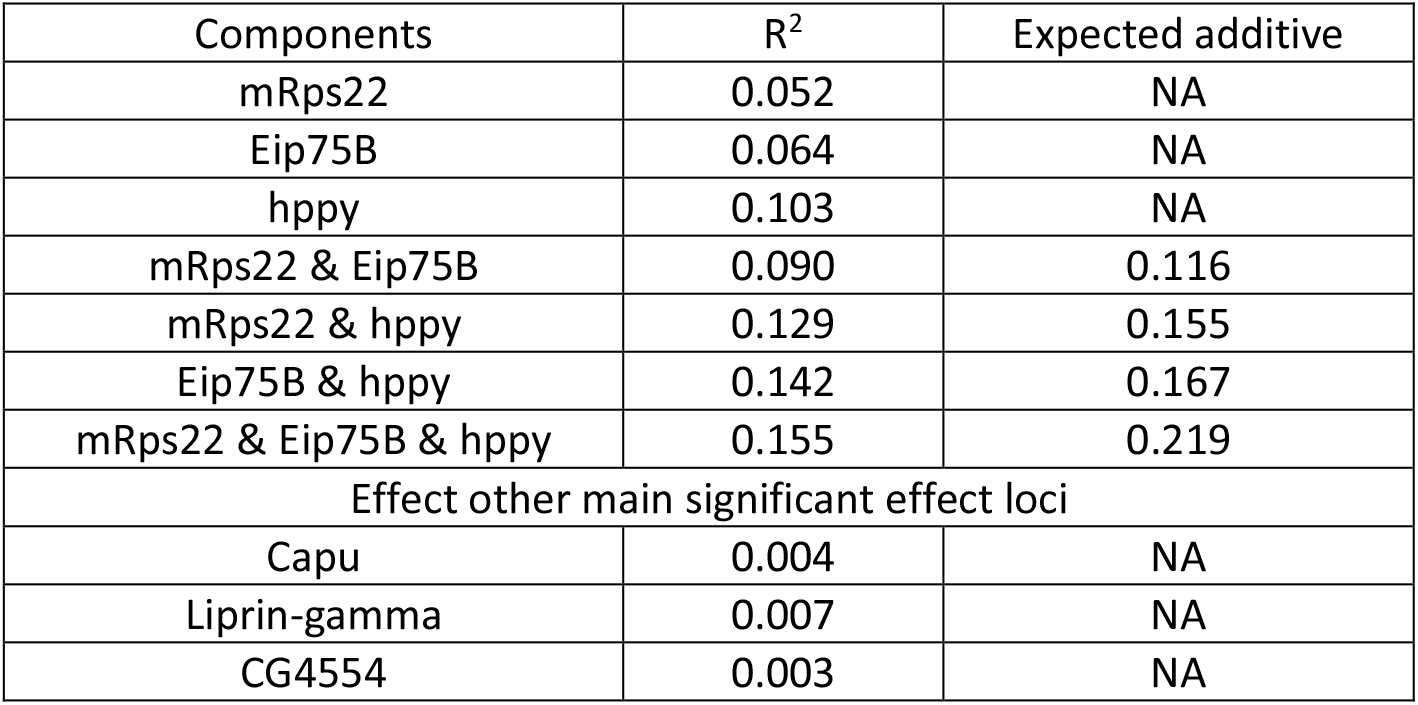
Variance components of fixed effects of major effect loci. The R^2^ is an estimation of the variance explained by a component. The expected additive effect is calculated as the sum of the single R^2^ values.

## Notes

### Competing Interest Statement

The authors have declared no competing interest.

## References

Barton, K. (2022). “MumIn: multi-model inference.” https://CRAN.R-project.org/package=MuMIn.

Barton, N. H., A. M. Etheridge and A. Veber (2017). “The infinitesimal model: Definition, derivation, and implications.” Theoretical Population Biology 118: 50–73.

Bates, D., M. Machler, B. M. Bolker and S. C. Walker (2015). “Fitting linear mixed-rffects models using lme4.” Journal of Statistical Software 67(1): 1–48.

Benjamini, Y. and Y. Hochberg (1995). “Controlling the false discovery rate - a practical and powerful approach to multiple testing.” Journal of the Royal Statistical Society Series B-Statistical Methodology 57(1): 289–300.

Bryk, B., K. Hahn, S. M. Cohen and A. A. Teleman (2010). “MAP4K3 regulates body size and metabolism in Drosophila.” Developmental Biology 344(1): 150–157.

Carnes, M. U., T. Campbell, W. Huang, D. G. Butler, M. A. Carbone, L. H. Duncan, S. V. Harbajan, E. M. King, K. R. Peterson, A. Weitzel, S. S. Zhou and T. F. C. Mackay (2015). “The genomic basis of postponed senescence in Drosophila melanogaster.” Plos One 10(9).

Chen, A. L., D. Tiosano, T. Guran, H. N. Baris, Y. Bayram, A. Mory, L. Shapiro-Kulnane, C. A. Hodges, Z.C. Akdemir, S. Turan, S. N. Jhangiani, F. van den Akker, C. L. Hoppel, H. K. Salz, J. R. Lupski and D. A. Buchner (2018). “Mutations in the mitochondrial ribosomal protein MRPS22 lead to primary ovarian insufficiency.” Human Molecular Genetics 27(11): 1913–1926.

Clancy, D. J., Kennington, J. (2001). “A simple method to achieve consistent larval density in bottle cultures.” Dros Inf Serv 84: 168–169.

Fabian, D. K., K. Garschall, P. Klepsatel, G. Santos-Matos, E. Sucena, M. Kapun, B. Lemaitre, C. Schlotterer, R. Arking and T. Flatt (2018). “Evolution of longevity improves immunity in Drosophila.” Evolution Letters 2(6): 567–579.

Hoedjes, K. M., Kostic, H., Flatt, T., Keller, L. (2023). A single nucleotide variant in the PPARgamma-homolog Eip75B affects fecundity in Drosophila.” Molecular Biology and Evolution 40(2):msad018

Hoedjes, K. M., J. van den Heuvel, M. Kapun, L. Keller, T. Flatt and B. J. Zwaan (2019). “Distinct genomic signals of lifespan and life history evolution in response to postponed reproduction and larval diet in Drosophila.” Evolution Letters 3(6): 598–609.

Huang, W., T. Campbell, M. A. Carbone, W. E. Jones, D. Unselt, R. R. H. Anholt and T. F. C. Mackay (2020). “Context-dependent genetic architecture of Drosophila life span.” Plos Biology 18(3).

Kiel, C., C. A. Nebauer, T. Strunz, S. Stelzl and B. H. F. Weber (2021). “Epistatic interactions of genetic loci associated with age-related macular degeneration.” Scientific Reports 11(1).

Kirkwood, T. B. L. (1977). “Evolution of aging.” Nature 270(5635): 301–304.

Lam, D., S. Shah, I. P. De Castro, S. H. Y. Loh and L. M. Martins (2010). “Drosophila happyhour modulates JNK-dependent apoptosis.” Cell Death & Disease 1.

Lehtovaara, A., H. Schielzeth, I. Flis and U. Friberg (2013). “Heritability of life span is largely sex limited in Drosophila.” American Naturalist 182(5): 653–665.

Leips, J. and T. F. C. Mackay (2002). “The complex genetic architecture of Drosophila life span.” Experimental Aging Research 28(4): 361–390.

Luckinbill, L. S., R. Arking, M. J. Clare, W. C. Cirocco and S. A. Buck (1984). “Selection for delayed senescence in Drosophila melanogaster.” Evolution 38(5): 996–1003.

Mackay, T. F. C. (2002). “The nature of quantitative genetic variation for Drosophila longevity.” Mechanisms of Ageing and Development 123(2-3): 95–104.

May, C. M., J. van den Heuvel, A. Doroszuk, K. M. Hoedjes, T. Flatt and B. J. Zwaan (2019). “Adaptation to developmental diet influences the response to selection on age at reproduction in the fruit fly.” Journal of Evolutionary Biology 32(5): 425–437.

Medawar, P. B. (1952). An Unsolved Problem in Biology. London, H.K. Lewis.

Nuzhdin, S. V., E. G. Pasyukova, C. L. Dilda, Z. B. Zeng and T. F. C. Mackay (1997). “Sex-specific quantitative trait loci affecting longevity in Drosophila melanogaster.” Proceedings of the National Academy of Sciences of the United States of America 94(18): 9734–9739.

Pallares, L. F., A. J. Lea, E. V. Filippova, P. Andolfatto and J. F. Ayroles (2022). “Dietary stress remodels the genetic architecture of lifespan variation in outbred Drosophila.” Nature Genetics 55: 123 – 129.

Parker, G. A., N. Kohn, A. Spirina, A. McMillen, W. Huang and T. F. C. Mackay (2020). “Genetic basis of increased lifespan and postponed senescence in Drosophila melanogaster.” G3-Genes Genomes Genetics 10(3): 1087–1098.

Partridge, L., N. Prowse and P. Pignatelli (1999). “Another set of responses and correlated responses to selection on age at reproduction in Drosophila melanogaster.” Proceedings of the Royal Society B-Biological Sciences 266(1416): 255–261.

R Core Team, (2019). R: A language and envrionment for statistical computing. Vienna, R foundation for statistical computing.

Remolina, S. C., P. L. Chang, J. Leips, S. V. Nuzhdin and K. A. Hughes (2012). “Genomic basis of aging and life-history evolution in Drosophila melanogaster.” Evolution 66(11): 3390–3403.

Roff, D.A. (1992). The evolution of life histories. New York, Chapman and Hall.

Roff D.A. and T.A. Mousseau (1987). “Quantitative genetics and fitness: lessons from Drosophila.” Heredity 58: 103–118.

Rose, M. R. (1984). “Laboratory evolution of postponed senescence in Drosophila melanogaster.” Evolution 38(5): 1004–1010.

Saada, A., A. Shaag, S. Amon, T. Dolfin, C. Miller, D. Fuchs-Telem, A. Lombes and O. Elpeleg (2007). “Antenatal mitochondrial disease caused by mitochondrial ribosomal protein (MRPS22) mutation.” Journal of Medical Genetics 44(12): 784–786.

Semagn, K., R. Babu, S. Hearne and M. Olsen (2014). “Single nucleotide polymorphism genotyping using Kompetitive Allele Specific PCR (KASP): overview of the technology and its application in crop improvement.” Molecular Breeding 33(1): 1–14.

Smits, P., A. Saada, S. B. Wortmann, A. J. Heister, M. Brink, R. Pfundt, C. Miller, D. Haas, R. Hantschmann, R. J. T. Rodenburg, J. A. M. Smeitink and L. P. van den Heuvel (2011). “Mutation in mitochondrial ribosomal protein MRPS22 leads to Cornelia de Lange-like phenotype, brain abnormalities and hypertrophic cardiomyopathy.” European Journal of Human Genetics 19(4): 394–399.

Stanley, P. D., E. Ng’oma, S. O’Day and E. G. King (2017). “Genetic dissection of nutrition-induced plasticity in insulin/insulin-like growth factor signaling and median life span in a Drosophila multiparent population.” Genetics 206(2): 587–602.

Stearns, S. C. (1992). The evolution of life histories. New York, Oxford University Press.

Stearns, S. C., M. Ackermann, M. Doebeli and M. Kaiser (2000). “Experimental evolution of aging, growth, and reproduction in fruitflies.” Proceedings of the National Academy of Sciences of the United States of America 97(7): 3309–3313.

Wang, H., D. A. Bennett, P. L. De Jager, Q. Y. Zhang and H. Y. Zhang (2021). “Genome-wide epistasis analysis for Alzheimer’s disease and implications for genetic risk prediction.” Alzheimers Research & Therapy 13(1).

Warnes, G. R. (2003). “The genetics package.” R News 3(1): 9–13.

Williams, G. C. (1957). “Pleiotropy, natural selection, and the evolution of senescence.” Evolution 11(4): 398–411.

Wong, H.W.S. and L. Holman (2023). “Pleiotropic fitness effects across sexes and ages in the Drosophila genome and transcriptome.” Evolution 77(12): 2642–2655

Yang, J. A., B. Benyamin, B. P. McEvoy, S. Gordon, A. K. Henders, D. R. Nyholt, P. A. Madden, A. C. Heath, N. G. Martin, G. W. Montgomery, M. E. Goddard and P. M. Visscher (2010). “Common SNPs explain a large proportion of the heritability for human height.” Nature Genetics 42(7): 565–U131.

Zwaan, B., R. Bijlsma and R. E. Hoekstra (1995). “Direct selection on life-span in Drosophila melanogaster.” Evolution 49(4): 649–659.

